# Anastrozole mediated modulation of mitochondrial activity by inhibition of mitochondrial permeability transition pore opening: An initial perspective

**DOI:** 10.1101/724872

**Authors:** Somesh Kumar, Subhajit Ghosh, Neha Choudhary, Mohammed Faruq, Prem kumar Inderganti, Vikram Singh, Ravindra kumar Saran, Haseena Sait, Seema Kapoor

## Abstract

**Background:** Mitochondrial permeability transmembrane pore [mPTP] plays a vital role in alteration of the structure and function of mitochondria. Cyclophillin D is a mitochondrial protein that regulates mPTP function and a known drug target for therapeutic studies involving mitochondria. While aromatase inhibitor’s role on mPTP has been previously studied, the role of anastrazole on mPTP is not completely elucidated.

**Methods:** The role of anastrozole in modulating the mPTP was evaluated by docking and molecular dynamics using human cyclophillin D data. Peripheral blood mononuclear cells [PBMCs] of patients with mitochondrial disorders and healthy controls were treated with anastrazole and evaluated for mean fluorescence by flow cytometer. Spectrophotometry was employed for total ATP level estimation.

**Findings:** Anastrozole – cyclophillin D complex is more stable when compared to cyclosporine A – cyclophillin D. Anastrozole performed better than cyclosporine in inhibiting mPTP pore. Additional effects included reduction in mitochondrial swelling and mitochondrial membrane depolarization, decreased super oxide generations, caspase 3 intrinsic activity and cellular apoptosis levels and increase in ATP levels.

**Interpretation:** These results highlights the potency of anastrozole as a promising agent in ameliorating the phenotype by inhibiting the opening of mPTP pore. However, larger functional studies are required to validate the efficacy of this molecule as a therapeutic agent in mitochondrial disorders.

## 1. INTRODUCTION

Mitochondrial disorders are a group of inherited disorders that arise when mitochondria fail to harvest sufficient energy for the body to function properly. Due to diverse genetic and clinical manifestations, the resultant complex phenotype often poses a diagnostic challenge. As a consequence of multiple steps involved in energy synthesis, amelioration of symptoms in these disorders is often difficult with a single point of action. No single compound till date is known to ameliorate the phenotype by acting on multiple steps that lead to mitochondrial dysfunction. This places greater emphasis on developing an alternative drug that can be considered for next generation mitochondrial modulating drugs.

Mitochondrial dysfunction ultimately results in increased matrix calcium. Calcium [Ca^2+^] overload across the mitochondrial membrane results in massive production of oxygen derived free radicals. This as a consequence triggers the opening of the mega transition pore ^1^, also known as mitochondrial permeability transition pore [mPTP] that was first described by Haworth and Hunter ^2^. It is a complex structure comprising of several protein domains such as voltage dependent anion channel [VDAC] in the outer membrane, adenine nucleotide translocator [ANT] in the inner membrane, Cyclophillin D [CypD] in the matrix, and several other molecules ^3^. Under normal conditions, mitochondrial matrix holds CypD and the mPTP remains closed. However its opening is finely regulated by the plethora of factors, including changes in ionic Ca^2+^, pH, reactive oxygen species [ROS], adenosine diphosphate/adenosine triphosphate [ADP/ATP] levels and the expression levels of B-cell lymphoma-2 [Bcl-2], an antiapoptotic protein ^4^.

The depolarization of the mitochondrial membrane potential [Δψm] changes the dynamics of the ATP synthase leading to reduced production of ATP. This accelerates the cascade leading to energy depletion, secondary to cellular apoptosis [Figure-1]. The distribution of Δψm thus could be considered as a potential prognostic factor which may assess the degree of tissue dysfunction or damage ^5^.

**Figure 1:**
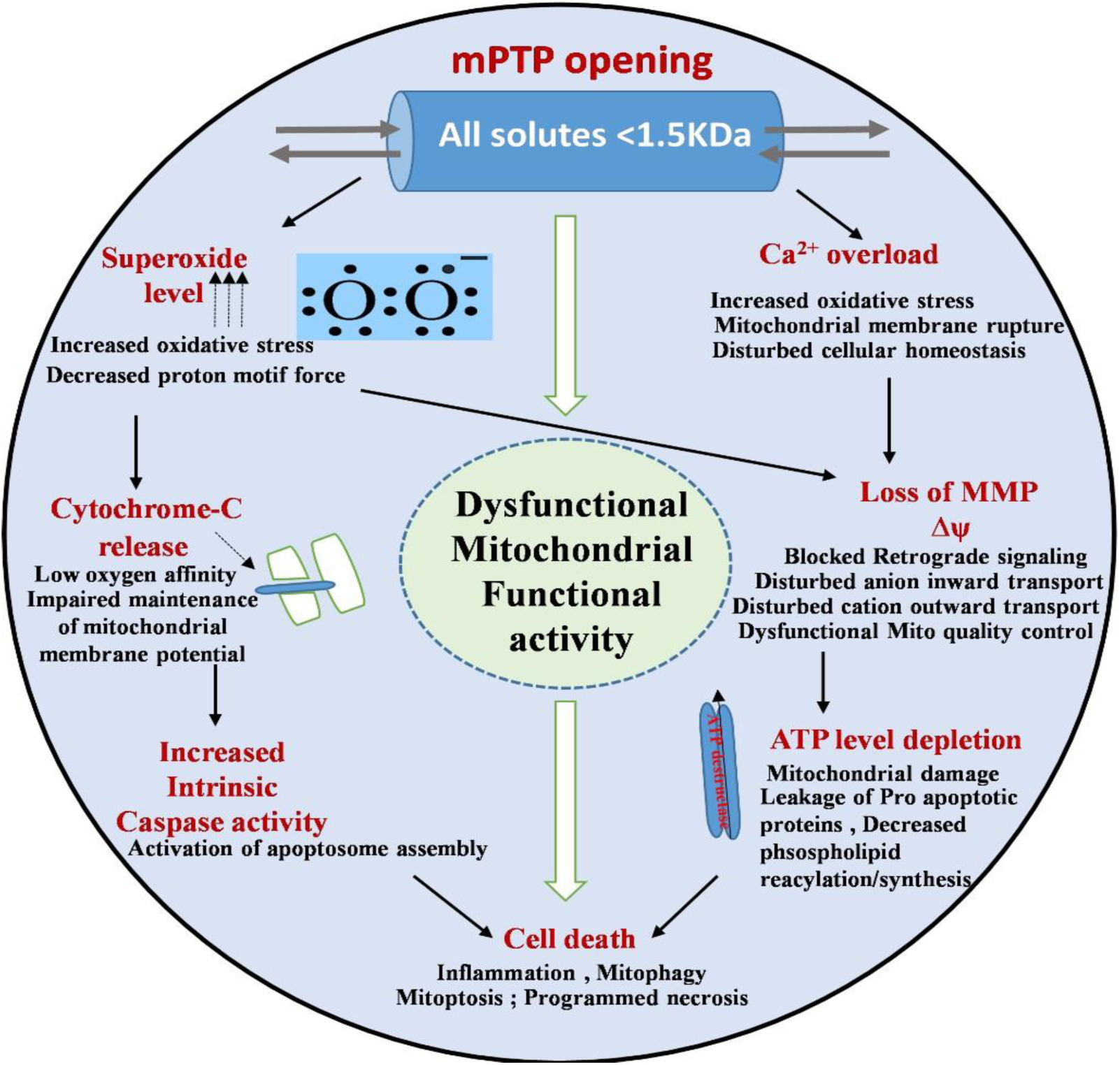
Cascades of mitochondrial dysfunctional events related to opening of mPTP pore: The mPTP pore opening allows the release of molecules of size up to 1500 Daltons leads to the cascades of unfavorable events such as mitochondrial swelling, rupture of outer mitochondrial membrane leading to release of cytochrome c oxidase into the cytosol, loss of mitochondrial membrane potential, depletion of ATP levels.

Precise targeting of mitochondria with that of compounds has been a long cherished dream of drug development team, pharmacologists, and researchers. Cyclosporine A [CysA], a potent immunosuppressant, inhibit calcineurin signaling by forming the complex with cyclophillin A [CypA] ^6–9^, which is a known mPTP blocker. Several others mPTP blockers targeting cyclophillin D with less trivial fashion includes non-immune-suppressant derivative of CsA, N-methyl-Val-4-cyclosporin A [MeValCsA] ^10^, Sanglifehrin A [SfA] ^11^, 2-aminoethoxydiphenyl borate [2-APB] ^12^, a non-immuno-suppressant agent, NIM811 [N-methyl-4-isoleucine cyclosporine] ^12^, anastrozole [a potent aromatase inhibitor] ^13^, local anesthetics, bongkrekic acid, N-acylethanolamines, butylhydroxytoluene, sulfhydryl reducing agents among many, carnitine, ADP, Sr^2+^, Mn^2+^, Mg^2+^ and H^+ 14^. Few organ specific supportive therapies for the alleviation of some symptoms ^15^ such as liver transplant ^16^ and allogenic haematopoietic stem cell transplantation ^17^ for neuro-gastrointestinal encephalopathy due to TYMP gene mutations, supplementation of N-acetylcysteine for ethylmalonic encephalopathy ^15^, Idebenone for patients with LHON ^18^, molecular bypass therapy aiming to restore deoxyribonucleoside triphosphate [dNTP] pools ^19,20^ for TK2 induced mitochondrial depletion syndrome, RRM2B gene induced nucleoside metabolism defect, Zinc finger nucleases [ZFN] ^21^ and transcription activator-like effector nucleases [TALENS] ^22^ have been experimentally used to manipulate the ratio of mutant and wild type of mitochondrial DNA [mtDNA] in cell lines.

Literature regarding the use of anastrozole as a selective and potent aromatase inhibitor suggests that it also controls mPTP opening ^13^. Loredano Moro et al in their study demonstrated the role of anastrozole on modulation of mPTP pore in aromatase knockout [ArKO] mice.

We hypothesized that opening of the mPTP is most likely to be linked to multi-complex unit assembly rather that an isolated function and its modulation reflects the change in mitochondrial activities. Hence, we decided to perform the two tier study in which we first evaluated [molecular docking] the orientation of anastrozole to cyclophillin D [a key regulator of mPTP] and the physical movement [molecular dynamics] simulation of its active molecules during the interaction with cyclophillin D. We also intend to evaluate the effects of anastrozole [aromatase inhibitor] on modulation of mPTP and its effects on the mitochondrial cascade such as the Δψm, superoxide generation, ATP levels, caspase 3 activity and apoptosis in the PBMCs.

## 2. RESULTS

10 Patient’s with mitochondrial disorders [Supplementary table S1 and figure S1 respectively] were recruited as per the E. morava et al. diagnostic criteria ^23^. PBMCs were isolated for evaluating mitochondrial downstream function.

### 2.1. Docking and MD simulation analysis of molecular interaction between Cyclosporine A-Cyclophilin D and Anastrozole - Cyclophilin D complexes

Based on the active-site residues of Cyclophilin D obtained from the available literature ^24^, protein-ligand molecular dockings were performed using AutoDock programs. To assess the accuracy and performance of docking procedure, the cyclosporine A docked structure was superimposed with its crystallized structure that resulted in a low RMSD value [∼0.5 Å]. The protein-ligand interacting interface of both the complexes for cyclosporine A *i.e.* one obtained from *in-silico* docking procedure and other from the crystallographic studies were also checked for amino acid residues mediating the interactions. The conformations of *in-silic*o docked and ligand-bound crystal structure were found to exhibit a conserved pattern of ligand binding in the active site. Several key residues involved in the binding pocket of cyclophillin D, including ARG55, PHE60, MET61, GLN63, GLY72, ASN102, ALA103 and HIS126 are shared at the level of inter-molecular interactions in both these complexes, indicating towards the reliability of the used docking protocol [Figure 2].

**Figure 2.**
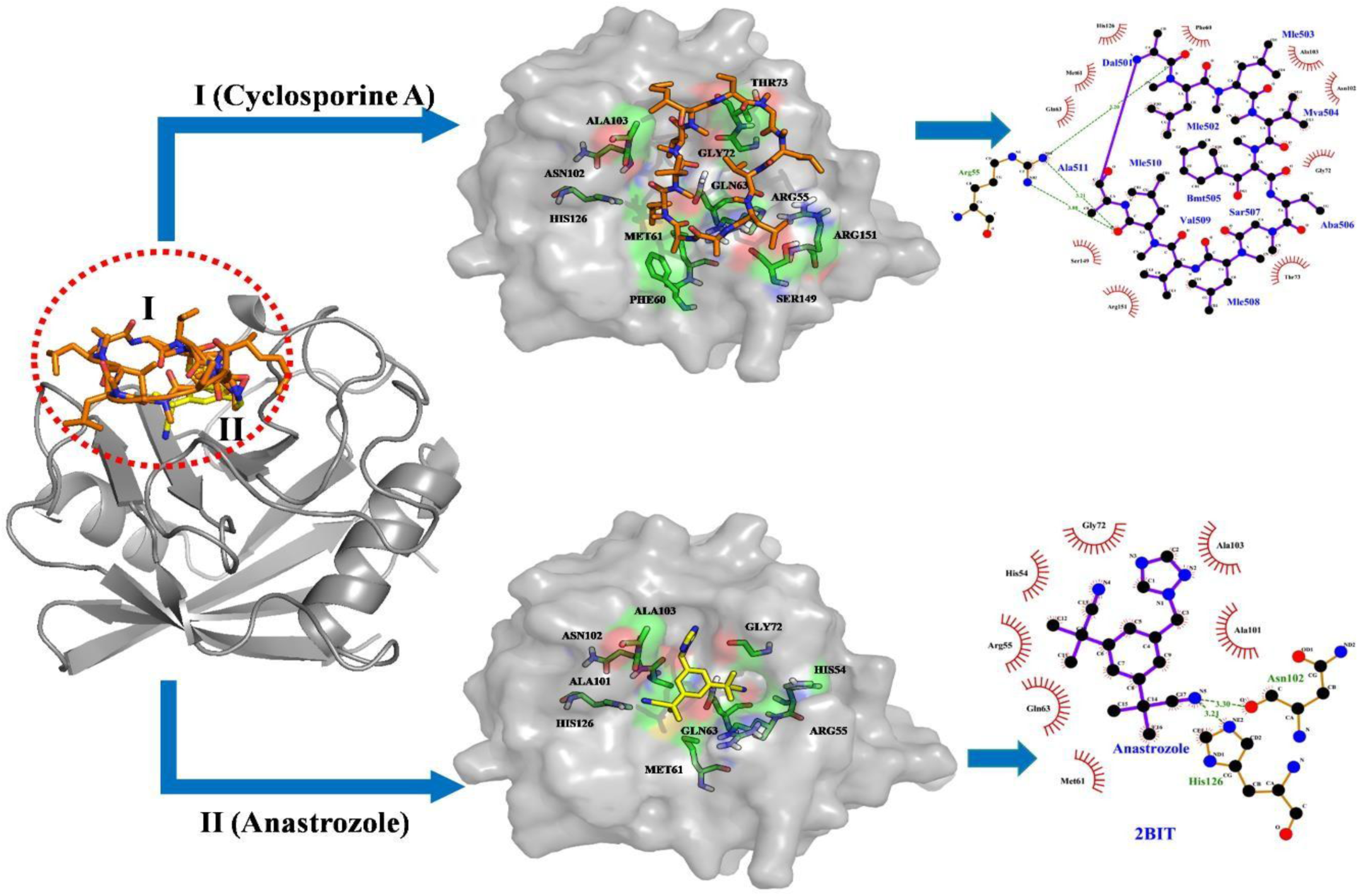
The docked complex of Cyclosporine A and anastrozole with Human cyclophillin D. The superimposed cartoon representation of cyclosporine A-cyclophillin D complex and anastrozole-cyclophillin D complex, showing the relative position of the docked cyclosporine A [represented as orange coloured sticks; I] and anastrozole [represented as yellow coloured sticks; II] inside the active site of protein [represented in the grey coloured ribbon layout]. Upper section represents cyclosporine A docked to the protein active site with the binding energy value of 5.0 kcal/mol. For anastrozole, the docking resulted in the binding energy value of −5.7 kcal/mol. The docked conformation and the protein residues interacting with anastrozole are represented in the lower panel. In the last terminals of both the sections are given the Ligplot+ analyses of docked complexes showing protein residues involved in hydrophobic interaction [represented in arcs] and hydrogen bonding [represented in dashed lines].

All the binding pocket residues of cyclophillin D are shown in Figure 3A. Using the similar docking protocol, anastrozole was also docked at the active site which resulted in the binding energy value of −5.7 kcal/mol compared to −5.0 for cyclosporine A. The docking results revealed that ligand [cyclosporine A] bound crystal structure and anastrozole docked complex share similar binding residues at the active site [ARG55, MET61, GLN63, GLY72, ASN102, ALA101, ALA103 and HIS126].

**Figure 3.**
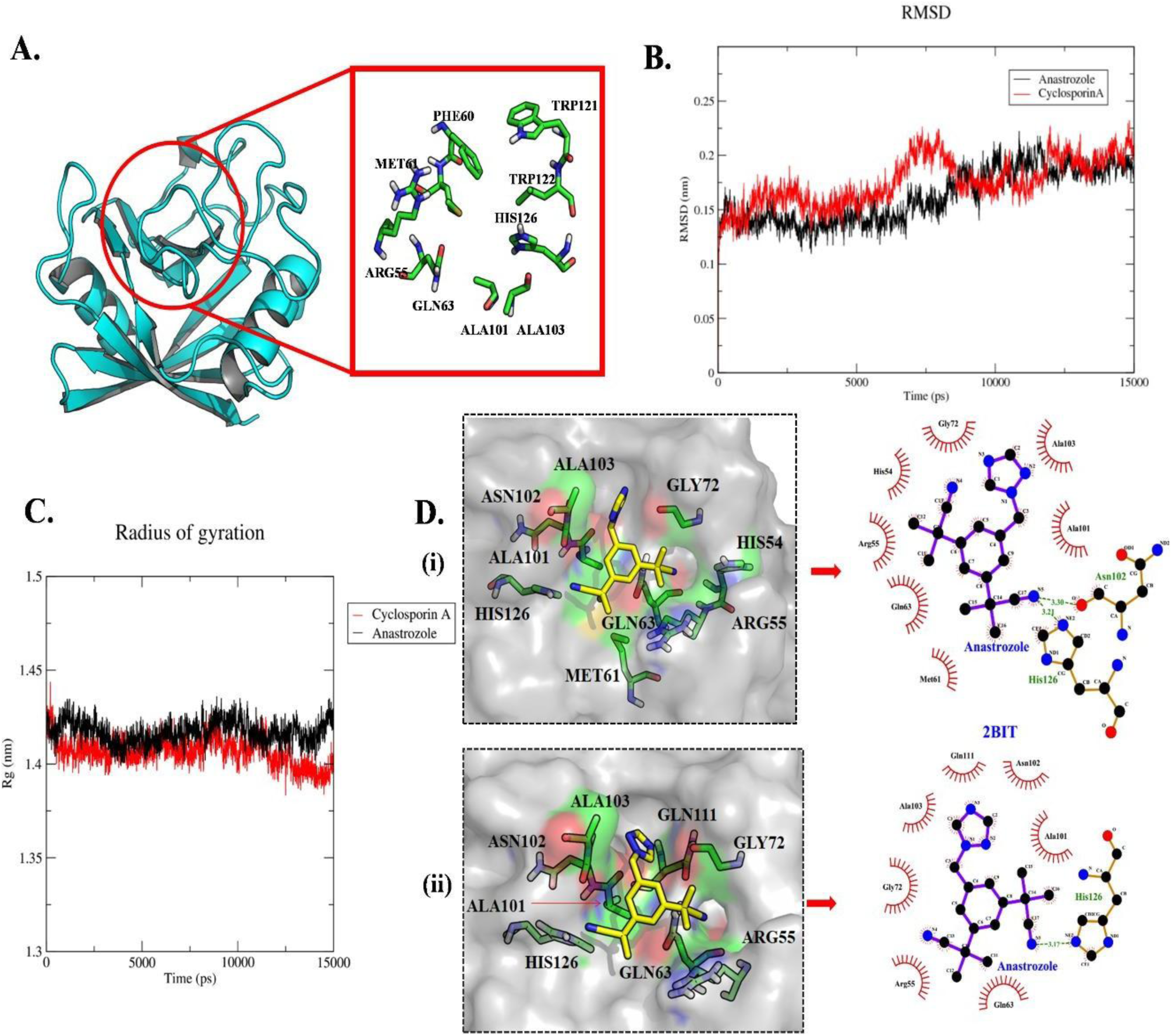
Molecular dynamics [MD] simulation showing stable interaction of Human Cyclophilin D with anastrozole. A) The tertiary structure of Human cyclophillin D [2BIT] with its active site residues highlighted in a box. The protein is depicted in cartoon representation [cyan colour] while active site residues are represented as green coloured sticks. B) Root mean square deviation [RMSDs] and C) radius of gyration [Rg] of Cα atoms during the 15ns of the MD simulation of anastrozole-cyclophillin D and cyclosporine A-cyclophillin D complexes. D) Anastrozole docked in the protein active site before and after molecular simulation showing hydrogen bonded and hydrophobic interacting residues of cyclophillin D (i) Before molecular simulation -- His126 and Asn102 form two H-bonds. His54, Arg55, Met61, Gln63, Gly72, Ala101 and Ala103 are involved in hydrophobic interactions, (ii) After molecular simulation -- His126 forms one H-bond. Arg55, Gln63, Gly72, Ala101, Asn102, Ala103 and Gln111 form hydrophobic contacts.

Additionally, we have found that anastrozole was involved in the formation of a hydrogen bond with the protein at HIS54, which further stabilizes anastrozole in the binding pocket [Figure 3A]. For confirming the reliability of anastrozole binding mode inside the protein active site, stability of anastrozole within the protein cavity was compared with that of the known inhibitor cyclosporine A by performing the molecular dynamics simulations of both the complexes [as detailed in the material and methods section]. The resulting simulation trajectories when analyzed for backbone stability [in terms of RMSD] were found to fluctuate around 0.15nm for both the ligands [as shown by red and black lines in Figure 3B]. For anastrozole the RMSD value range was 0.005 to 0.222nm, while for cyclosporine A it was 0.004 to 0.232nm. The trajectories of radius of gyration [R_g_] with respect to time varied between 1.38 to 1.44 Å for anastrozole and between 1.39 to 1.43 Å for Cyclosporin A respectively [Figure 3C]. These values were showing increased fluctuations in the early phases of simulation, but a sustained plateau was obtained as the simulation proceeded, indicating the stability of conformation. For the comparative analysis, ligand binding conformations before and after the simulation was generated and studied in detail [Figure 3D]. ARG55, GLN63, GLY72, ALA101 and ASN102, ALA103, GLN111 and HIS126 formed strong hydrophobic contacts with the anastrozole. However, the nitrile group of anastrozole moved away from MET61 and ASN102, breaking the H-bond and making new hydrophobic interactions. Interestingly, this led the imidazole ring to move deeper into the binding pocket and form new hydrophobic contact with GLN111; an active site residue [Figure 3D]. Also, the H-bond between the HIS126 and anastrozole got much stronger [from 3.30 Å to 3.17 Å]. Anastrozole-cyclophillin D binding interface was also checked for the involvement of Arg55, a key pocket residue referred as catalytic arginine for the cyclophillin family ^25^. The sustained interaction of anastrozole with Arg55, during the MD simulation represents the ability of ligand to facilitate reaction pathway.

### 2.2. Anastrozole’s role in the inhibition of mPTP pore opening

The mPTP opening in PBMCs was determined on the basis of change in green fluorescence intensity in the presence of and absence of H_2_O_2_ [Figure 4]. The decrease in fluorescence using CoCl_2_ was greater in case group than the control group (p=0.006). Addition of H_2_O_2_ caused a further significant decrease in green fluorescence in the case group (p<0.001) indicating H_2_O_2_ induced opening of mPTP. While PBMCs exposed to H_2_O_2_ was treated with the drug at its nontoxic concentration [Supplementary figure S2], there was a significant increase in the green fluorescence in case group (p=0.001) [Figure 4C]. These results were more pronounced in the case group when compared to control group. While H_2_O_2_ induced early and late apoptosis in both case and control group after 6 hours of incubation, anastrozole significantly reduced the H_2_O_2_ induced apoptosis in both the groups (p=0.001).

**Figure 4:**
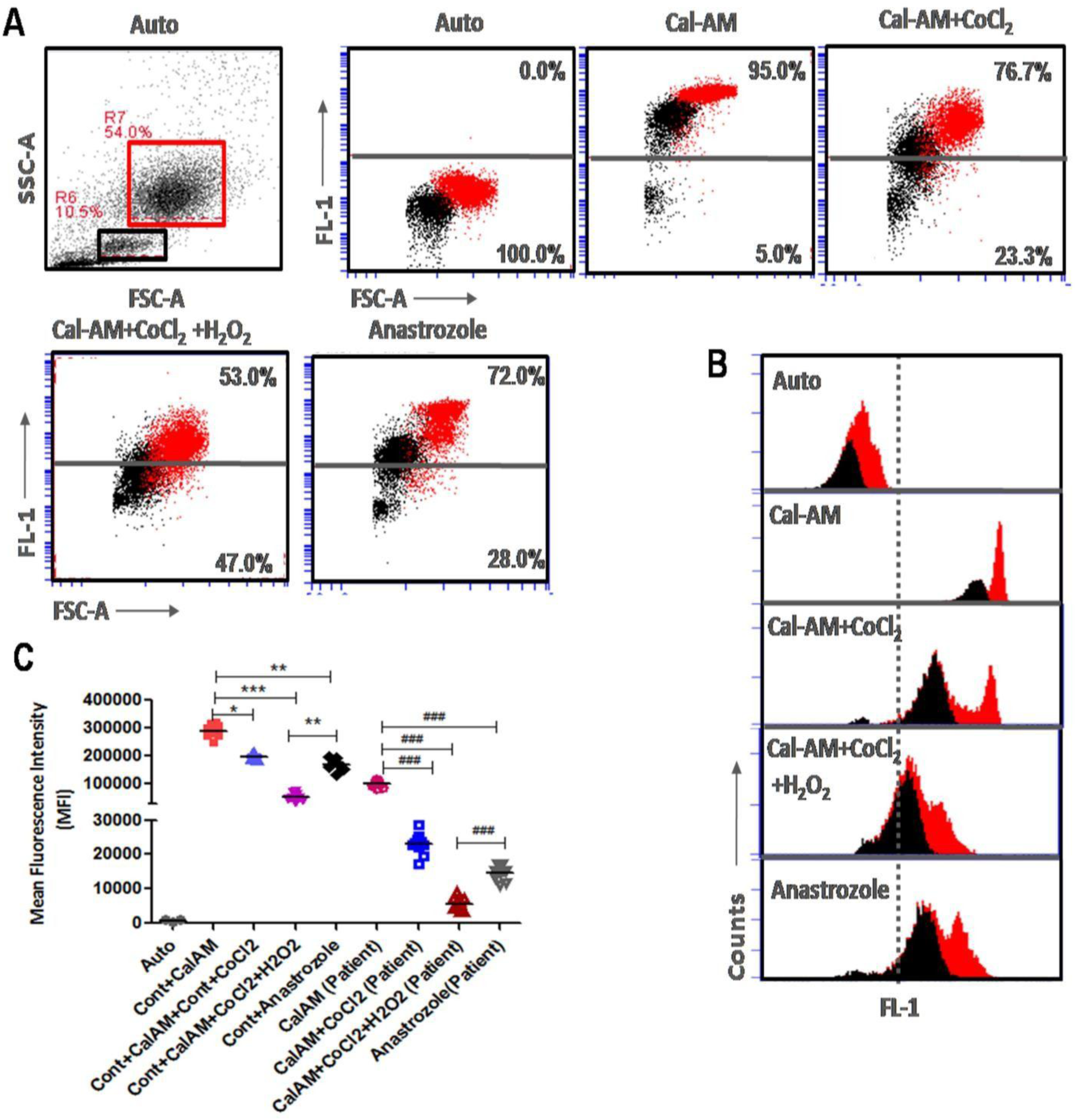
Anastrozole mediated modulation on mitochondrial transmembrane pore opening. Panel **A** and **B** represents gating strategy for flowcytometry analysis of lymphocyte [black square] and granulocyte [red square] in PBMCs and their changes in fluorescence upon different treatments. Panel **C** shows anastrozole inhibited H_2_O_2_ induced calcium retention capacity decrease in mean fluorescence intensity. Data represents as mean ± SEM [n = 10 subjects; control and patient respectively] and comparisons were done for statistical significance using unpaired t-test (*p<0.05, **p<0.01, **###**/***p<0.001).

### 2.3. Anastrozole modulated mitochondrial membrane potential

CCCP significantly reduced MMP and the fluorescence of preloaded Cal-AM in healthy (p=0.001) and patient’s PBMCs (p=0.001) in comparison to their untreated controls [Figure 5A]. However, treatment with anastrozole significantly reverted this CCCP-induced changes in control PBMCs (p=0.001) and in patient’s PBMCs (p=0.001).

**Figure 5:**
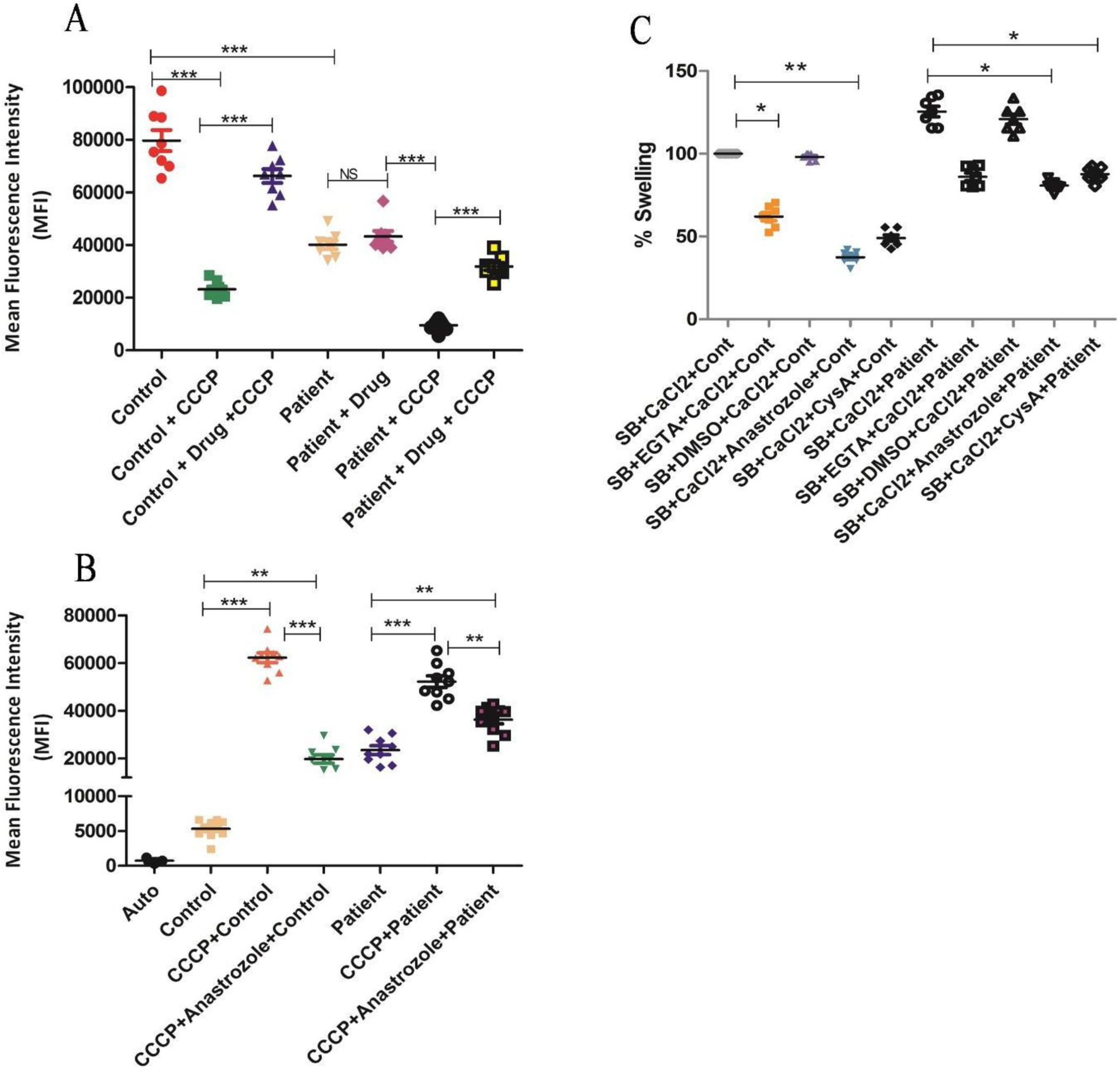
Quantitative modulation of anastrozole mediated different mitochondrial parameters. Changes in mitochondrial membrane potentials (**A**), superoxide level **(B)** and mitochondrial swelling **(C)** Estimated after treatment with anastrozole in healthy and patient PBMCs. Data represents as mean ± SEM [n = 10 subjects; control and patient respectively] and comparisons were done as indicated using unpaired t-test (*p<0.05, **p<0.01, ***p<0.001).

### 2.4. Anastrozole modulated superoxide generation

Superoxide generation was higher in case group than the control group (p<0.001). However on treatment with, there was significant increase in superoxide in control group [Figure 5B]. Addition of anastrozole to CCCP treated mitochondria caused a significant reduction in the superoxide generation, more in the case group in comparison to the control group (p=0.002).

### 2.5. Anastrozole’s role in calcium induced mitochondrial swelling

The patient’s mitochondria underwent rapid swelling in the presence of calcium, whereas anastrozole significantly inhibited calcium induced swelling [Figure 5C]. Cyclosporin A also significantly prevented swelling in patients’ mitochondria. However, while this data was compared with healthy subjects, a significant difference was observed

### 3.6. Anastrozole’s role in preventing loss in ATP levels and controlled apoptosis

CCCP reduced the total ATP level in comparison to the untreated control (p=0.003) [Figure 6A], whereas, drug treatment significantly ameliorated these changes (p<0.001). Further, there is an increase of cell death observed in patients PBMCs and suppressed by drug treatment while compared with healthy PBMCs [Figure-6B, p<0.001]. However, drug shown the similar results in CCCP treated PBMCs [Figure-6B, p<0.001]. To support these results, we evaluated caspase-3 level in both PBMCs and observed an elevated level of caspase-3 in patient PBMCs [Figure-6C, p<0.001]. Treatment with anastrozole significantly suppressed caspase-3 level in patient PBMCS and CCCP induced groups as compared with healthy and untreated group [Figure-6C, p<0.01].

**Figure 6:**
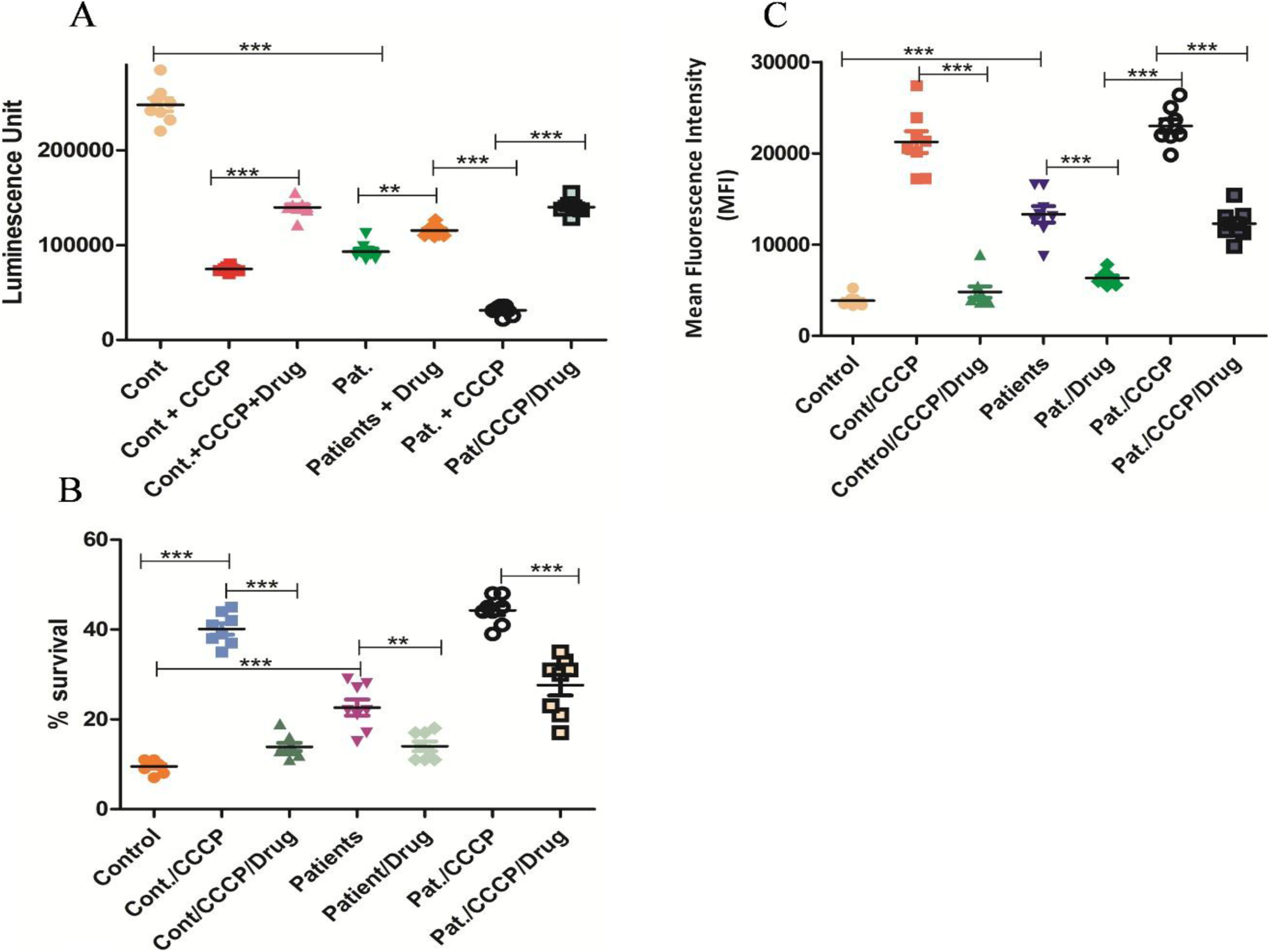
Effect of anastrozole on mitochondrial energy production and cell death. **(A)** Anastrozole inhibited the reduction of total ATP level after treatment with mitochondrial proton gradient uncoupling agent CCCP. (**B** and **C**) The quantified value of H_2_O_2_ induced apoptotic fraction [%annexin V positive cells expressed as %apoptosis] and caspase −3 levels in healthy and patient PBMCs which treated with anastrozole. Data represents as mean ± SEM [n = 10 subjects; control and patient respectively] and comparisons were done as indicated using unpaired t-test (*p<0.05, **p<0.01, ***p<0.001).

## 4. DISCUSSION

Mitochondriopathies accounts for majority of the common neurometabolic and neuromuscular disorders. No definitive therapy is available that directly modulates the function of mitochondria either by increasing the proportion of healthy mitochondria or by modulating the effects of mutated variants. The effects of various small molecules in mitochondrial disorders are currently being investigated in clinical and preclinical studies ^26^. The current study highlights the three new observations: (1) Anastrozole is found to be relatively more stable on interaction with cyclophillin D with few conformational changes in comparison to the known Cyclosporine A-Cyclophillin D complex; (2) Mutation in mitochondrial governed genes leads to unfavorable modulation in mitochondrial activity and penultimately leads to cell death by an opening of mPTP; (3) Anastrozole modulates the mutant mitochondrial function by inhibiting mPTP opening and reducing the cell death in PBMCs of patients with mitochondrial disorder. Using literature survey, the binding pocket residues of the protein were identified. Significant interactions of anastrozole at these binding pocket residues were obtained by the docking studies. Identification of common interacting residues among the docked anastrozole and cyclosporine A bound crystal structure of cyclophillin D showed that the binding potential of anastrozole is noteworthy. Also, H-bond integration of the anastrozole with His-126 might be playing a crucial role in stabilizing the protein-ligand complex [Figure 3D]. Further, binding affinity of the anastrozole was found to be relatively stable compared to its known inhibitor [cyclosporine A] as suggested by the obtained biding energy values. The structural fluctuations in the docked protein-ligand complexes were also accessed using molecular simulations studies. RMSD values calculated over the simulation of 15 ns for anastrozole suggested that the compound can form stable protein-ligand complex with cyclophillin D. RMSD fluctuations in cyclophillin D complex with the test molecule [anastrozole] and control ligand [cyclosporine A] were found to be comparable with each other, an observation that further supports the good inhibitory potential of Anastrozole [Figure 3B]. Monitoring of R_g_ pattern depicted compactness and less variation in the Anastrozole-Cyclophillin D complex as compared to the control system [shown in Figure 3C]. Further, hydrophobic interactions mediated by the catalytic arginine [Arg-55] could be suggested as a driving factor for the ligand efficacy of anastrozole towards cyclophillin D. Most of the interactions revealed by the docking studies were retained in the MD simulation. No significant drift in the MD simulation of anastrozole shows that the compound possesses the ability to form a stable protein-ligand complex. This study also highlights that mitochondria of patient’s PBMCs had high calcium overload [Figure 5C] and the uncontrolled opening of the mPTP leading to cell death [Figure 6B]. The opening of mPTP occurred as a response to an apoptotic stimulus, resulting in the release of different apoptogenic molecules ^27^. Several factors increased the sensitivity of conformity changes in mitochondria to eventually forcing the opening of mPTP, including increased ROS generation, depletion of ATP levels and depolarization of mitochondrial membrane potentials ^28,29^. In our study, we also observed the elevated mitochondrial superoxide production [Figure 5B], depolarized membrane [Figure 5A] and low ATP level [Figure 6A] in PBMCs of the patients. It was also reported that the loss of MMP and superoxide production during a stress condition, induces cell death ^30^. Studies have also suggested that mutation in the mitochondrial genome affected the electron transport chain activity and thereby ATP production. Till date, no single FDA approved molecule is available for the treatment of mitochondrial disorders ^31^. However, several drug targets were reported for these disorders ^32,33^ including mitochondrial membrane protein known as cyclophillin-D, that regulates the mitochondrial transmembrane pore. Aromatase enzyme also regulates mPTP opening ^34–36^. However, few reports showed that deficiency of aromatase enzyme inhibits the mPTP opening ^13^. In the same study ^13^, authors didn’t observe any significant change in the membrane depolarization, ROS generation and ATP level depletion in the liver mitochondria of genetically knockout [aromatase] mice. However, in our study, we observed that anastrozole restored mitochondrial membrane potential [Figure 5A], reduces the superoxide level [Figure 6B] and cell death [Figure 6B], and this could be due to the temporary closing of mPTP by the drug. These results are also in line with the modulation capability of the known mPTP inhibitor cyclosporine A.

Anastrozole seems to be a promising agent in ameliorating the phenotype by regulating the activity across the mPTP pore. Larger functional studies including the levels of cytochrome c oxidase may give insight into mechanistic modulation of the mPTP pore carried out by this agent. Clinical trials with this molecule may therefore provide insight into its advantages, selectivity and dosage regulation.

## 5. MATERIALS AND METHODS

The study carried out with the necessary approval from the Maulana azad medical college affiliated Institutional Ethical Committee (MAMC-IEC) board and complies all necessary steps with the Declaration of Helsinki. The cross-sectional study adhered to the Ethical Principles for Medical Research Involving Human Subjects, World Medical Association Declaration of Helsinki. A convenient patient sample size of 10 was taken due to difficulty in diagnosing the disorder into the consideration along with 10 healthy control samples. Informed consent was obtained from all subjects.

### 5.1. Isolation of PBMCs

Ten healthy control [age-sex matched] and ten patients with mitochondrial disorders were selected by using modified E Morava et al ^23^ disease scoring system. 2 ml whole blood from each group [controls and patients] were collected in heparinized vials [BD biosciences]. Red blood cells [RBC’s] was lysed, using 1x ammonium chloride potassium [ACK] solution and washed 2 times with 1x phosphate buffer saline [PBS, pH 7.2]. For accurate volumetric measurements, fluid calibration in BD Accuri C6-flow cytometer, was done on the day of counting, following the manufacturer’s recommendations. The absolute volumetric count was calculated as follows: Total cell count [cells/μL] = [no. of viable cells of interest/ measured sample volume] × dilution factor.

### 5.2. Drug toxicity assay

The appropriate dosage selection of anastrozole was identified by using the methylthiazol tetrazolium [MTT] assay on human embryonic kidney cells [HEKs]. HEKs were treated with different concentrations of the drug, in RPMI 1640 media for 6 hours in CO_2_ incubator. After removing the media, the obtained formazan crystals were stabilized by the addition of dimethyl sulfoxide [DMSO]. Absorbance was measured [570 nm] using a spectrophotometer and a plate reader, and growth inhibition was calculated.

### 5.3. DOCKING AND MOLECULAR SIMULATION STUDY

#### 5.3.1. Preparation of cyclophillin D and ligand for molecular docking

Three-dimensional crystal structure of Cyclophilin D [PDB ID: 2BIT] resolved at 1.7Å ^37^ was obtained from protein data bank in order to use it as a docking template. Cyclosporin A, a known Cyclophilin D receptor antagonist was selected as the positive and/or reference compound to compare it with target compound anastrozole. These two compounds were prepared for molecular docking, processed and minimized with the help of Open Babel version 2.4.1 ^38^.

#### 5.3.2. Parameter optimization and molecular docking

AutoDock version 4.2 was used to perform the docking studies mainly due to its processing speed and superior ability for binding mode prediction ^39^. First of all, information about the binding pocket of Cyclosporine A in Cyclophilin D was obtained from available literature ^24^. Optimal grid structure was selected and coordinates covering the whole binding site of the Cyclophilin D were chosen for further studies. The parameters adopted were, center_x 24.28 Å, center_y 32.51 Å and center_z 26.04 Å with the grid size size_x 25 Å, size_y 25 Å and size_z 25Å. Both control and test ligands were docked in the binding pocket of the Cyclophilin D and comparisons were made between the conformations of protein complex docked with test ligand [anastrozole] and control ligand [Cyclosporine A] bound crystal structure available in PDB ^24^. The best binding orientations of both the ligands [*i.e.* the orientations with minimum binding energy] within the protein active site were estimated based on binding energy values.

#### 5.3.3. Molecular Dynamics [MD] simulations study

MD was conducted using GROningen Machine for Chemical Simulation 5.0 [GROMACS 5.0] ^40^ using the GROMOS96 53a6 ^24^ force field. Topologies of the ligands were obtained from PRODRG server ^41^. To avoid steric hindrance and to obtain stable conformation, energy minimization of the system was performed using steepest descent algorithm. Simple Point Charge [SPC] model was applied to solvate the system ^42^ and four Cl-counter ions were added to satiate the electro-neutrality condition. Parrinello-Rahman pressure coupling ^43^ and V-rescale temperature coupling ^44^ using coupling constant of 2.0 for pressure and 1.0ps for temperature were used to stabilize the system. Particle Mesh-Ewald [PME] method ^45^ for determining the long-range electrostatic interaction was employed and the LInear Constraint Solver [LINCS] ^46^ algorithm was used to constrain all the bonds. The complexes in the designed medium were equilibrated for 100ps [1 bar pressure and 300K temperature] in NVT and NPT ensemble. After equilibration, 15 ns MD simulation was performed on the system and the trajectories were analyzed for root mean square deviation [RMSD] and radius of the gyration [Rg]. Graphical Advanced Computation and Exploration [GRACE] utility of the GROMACS package was used to generate and analyze the graphical plots. Ligplot+, a graphical system that utilizes 3D coordinates to generate the interaction maps was used to visualize the interaction present in protein-ligand complex ^47^. Molecular graphics were generated using PyMOL ^48^.

### 5.4. CASCADES OF EVENTS IN MITOCHONDRIA

#### 5.4.1. Mitochondrial swelling assay and drug treatment

Mitochondrial swelling assay was detected as described earlier. Mitochondria were suspended in a swelling assay buffer [SB; 150 mM sucrose, 50 mM potassium chloride, 2 mM dipotassium phosphate, 5 mM succinic acid, 20 mM Tris-HCl, pH7.4]. As cyclosporine A is well known mPTP blocker, we planned to evaluate the efficacy of anastrozole by comparing with cyclosporine A in both case-control group. Cyclosporine A and Anastrozole [2.5uM respectively] was incubated for ten minutes and then, 100 μM calcium chloride was added and O.D. [540 nm] was measured per minute for 30 minutes by using a spectrophotometer.

#### 5.4.2. Mitochondrial permeability transition pore [mPTP] assay

To evaluate the mPTP openings in PBMCs were assessed using the Mito Probe Transition Pore Assay Kit [Molecular Probes] according to the manufacturer’s instructions with slight modifications. Briefly, drug-treated cells were loaded with 50 nM of calcein-acetoxy methyl ester [Cal-AM] at 37 °C for 20 minutes. Cells were then washed with Hanks’ balanced salt solution buffer twice. Then, cells were treated with 0.4 mM Cobalt chloride [CoCl2] or 0.4 mM CoCl_2_ plus 1 mM hydrogen peroxide [H_2_O_2_] for 10 min at 37°C. The samples were then analyzed using a BD Accuri C6 flow cytometer with appropriate excitation and emission filters for FITC. The green fluorescent dye calcein acetoxy methyl [Cal-AM] ester get de-esterified by esterase enzyme after entering the cytoplasm and mitochondria. CoCl_2_ quenches the fluorescence in the cytoplasm but not in mitochondria, due to its non-permeable nature towards the mitochondrial membrane, whereas H_2_O_2_ indices mPTP pore opening and CoCl_2_ could easily enter the mitochondria and reduces the fluorescence. The change in fluorescence intensity between samples incubated with only Cal-AM/CoCl_2_ and samples incubated with Cal-AM/CoCl_2_+H_2_O_2_ indicates mPTP-induced opening by H_2_O_2_.

#### 5.4.3. Mitochondrial superoxide generation assay

To determine mitochondrial superoxide production in healthy and patients PBMCs, dihydroxy ethidium staining was performed. Isolated PBMCs were treated with the drug for ten minutes and then incubated with H_2_O_2_. After washing, cells were treated with 1µM of dihydroethidium and immediately analyzed using BD Accuri C6 software.

#### 5.4.4. Mitochondrial membrane potentials assay

PBMCs were incubated with the drug for 10 minutes in a CO_2_ incubator and then treated with H_2_O_2_ and/or protonophore-m-chlorophenylhydrazone [CCCP] for another 30 minutes. Cells were washed twice and stained with 10 nM Rhodamine 123 [Rh123] and analyzed using BD Accuri C6 software.

#### 5.4.5. ATP assay

The cellular total ATP level quantification was done after completion of treatment using a commercially available chemilluminescence assay kit [Sigma Aldrich, USA]. Protocol was followed according to manufacturer instructions.

#### 5.4.6. Annexin-V/PI and caspase-3 assays

Cell death was quantified using Annexin-V/PI kit and protocol was followed according to manufacturer’s instructions. Briefly, after completion of treatment, cells were subjected to a binding buffer containing annexin V and propidium iodide and incubated at room temperature for 15 minutes and acquired by flowcytometry. Similarly, cleaved caspase-3 levels in isolated healthy and patient PBMCs was measured by following the kit manufacture’s instruction [Caspase-3 assay kit, Abcam, ab39383].

### 5.5. Statistical analysis

All the results are presented as mean ± standard error of the mean [SEM]. All statistical analysis were performed using GraphPad Prism from GraphPad Software [v5.0, La Jolla, CA, USA]. One way ANOVA repeated measure followed by boneferoni post-test were used for multiple comparisons. Student’s t-test was used for single comparison of analytes. p-value less than 0.05 were considered statistically significant.

## Acknowledgements

All the authors are grateful to Dr. Bidhan Chandra Koner [Department of Biochemistry, MAMC] and [Dr. Arun Kumar, DRDO-INMAS] for their technical support. All authors greatly acknowledge the organizational and infrastructural supports of the participating institutions.

## Author Contributions

Study conception and design – SK, SK, SG.

Data acquisition – SK, SG, NC.

Data Analysis – SK, SG, NC

Interpretation of Data – SK, SG, NC, MF, VS, PKI, RKS, HS, SK

Writing and/or revision – SK, SG, NC, HS, MF, RKS, VS, SK

Study Supervision – SK, MF, VS, RKS.

Study Administration – SK, SG, NC, SK, MF, VS, RKS, HS, PKI.

SK – Dr. Seema Kapoor (M.D)

SK – Somesh Kumar (PhD Scholar)

SG – Dr Subhajit Ghosh (PhD)

NC – Neha Choudhary (PhD. Scholar)

MF – Dr. Mohammed Faruq (PhD)

VS – Dr. Vikram Singh (PhD)

PKI – Dr. Prem Kumar Inderganti(PhD)

RKS – Dr Ravindra kumar Saran (M.D)

HS – Dr Haseena Sait (M.D)

