## Supplementary File for "Anastrozole mediated modulation of mitochondrial activity by inhibition of mitochondrial permeability transition pore opening: An initial perspective"

### SUPPLEMENTARY FILES:-

| S.No. | CLINICAL FINDINGS | MOLECULAR FINDINGS |
|---|---|---|
| **P-1** | 12 years male born of a non-consanguineous marriage, presented with clinical indications of gradual vision loss, neurological issues, murmuring and seizures. Muscle biopsy – No significant changes. | MT-ND4<br>c.1204G>A.(p.Val402Ile)<br>(Homoplasmic) |
| **P-2** | 5 year old male child born of a non-consanguineous marriage, presenting with clinical indications of gross motor delay, seizures, epileptic encephalopathy, hypotonia, dystonia, failure to thrive. | NDUFV1<br>Exon 8. c.1156C>T; (p.Arg386Cys)<br>(Homozygous) |
| **P-3** | 15 years old female born of a non-consanguineous marriage, presented with clinical indication of having progressive opthalmoplegia, with drooping of eyes since last 8-9 years. Overall dull vision. Muscle biopsy – COX- stain negative with SDH positivity. | MT-ATP6<br>c.158C>T; (p.Thr53Ile)<br>(Homoplasmic) |
| **P-4** | 20 years old female born of a non-consanguineous, presented with clinical indications of gradual vision loss and mild intellectual disability. Muscle biopsy – No significant changes. | MT-ND4<br>c.1204G>A; (p.Val402Ile)<br>(Homoplasmic) |
| **P-5** | 15 years old male born of a non-consanguineous marriage, presented with a clinical indication of painless loss of vision (past 12 months). | MT-ATP6<br>c.158C>T; (p.Thr53Ile)<br>(Homoplasmic) |
| **P-6** | 15 years old female born of a non-consanguineous, presented with clinical indications of gradual vision loss and mild intellectual disability. Muscle biopsy – No significant changes. | MT-ND4<br>c.1204G>A; (p.Val402Ile)<br>(Homoplasmic) |
| P-7 | 27 years old female born of a non-consanguineous marriage, presented with clinical indications of gradually progressive, proximal myopathy, proximal and distal muscle weakness with encephalopathy and inability to get up from sitting position. Muscle biopsy – Occasional COX negative fibres. | MT-ATP6<br>c.31G>A; (p.Ala11Thr)<br>(Homoplasmic) |
| **P-8** | 10 years old male born of a non -consanguineous, presented with clinical indications B/Loss of vision, with MRI – Orbit – indicating left optic nerve in the retro bulbar segment shows short segment thinning with no definitive evidence of any altered signal intensities. Muscle biopsy – No significant changes. | MT-ND4<br>c.1204G>A; (p.Val402Ile)<br>(Homoplasmic) |
| **P-9** | 2 years old male born of a non -consanguineous, presented with clinical indications seizures, encephalopathy with dystonia , failure to thrive | NDUFAF2<br>Exon.4 c.282_283delAC<br>(p.His94GlnfsTer13)<br>(Heterozygous) |
| **P-10** | 3 years old male child, born of a non-consanguineous marriage, presented with clinical indications of progressive lower limb muscle weakness. MRS shows a double lactate peak. | NDUFV1<br>Exon 8 c.1156C>T; (p.Arg386Cys)<br>(Homozygous) |

Supplementary table – S1: Clinical, mutational and histopathological findings in patients with mitochondrial disorders.

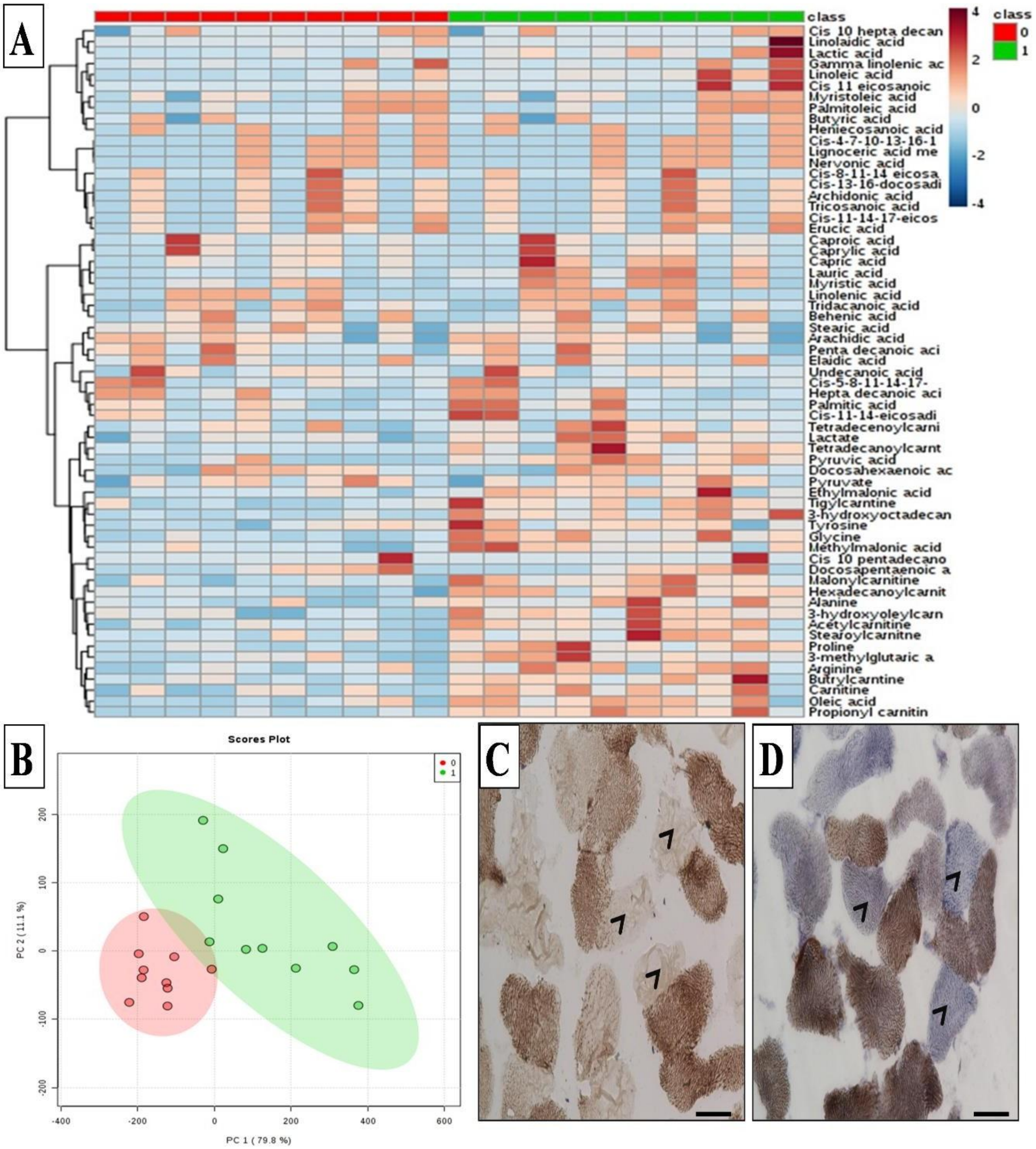


Supplementary Figure – S1**: Metabolic and Immnuo-histopathological profiling. (A)** The levels of 63 metabolites were z-score transformed to facilitate data representation and then plotted in a heat map. Each row corresponds to a unique metabolite, and each column corresponds to a biological replicate. (0 – Controls; 1 - Patients). Brown indicates high abundance, while blue shows low abundance. **(B)** Principle component analysis of selected data matrix of metabolites. Similarities and differences in metabolite abundances due to mutation in patient drove the clustering and separation of value within and healthy subjects respectively [0 – Controls (Red) ; [1 – Patients(Green)]. **In Pathological findings (C)** 40X Cox stain (Cytochrome oxidase) enzyme histochemistry- arrow reveal cox negative fibers **(P-7)**. **(D)** 40X Cox with SDH (succinate dehydrogenase) enzyme histochemistry- highlights the Cox negative fibers by only SDH stains **(P-3)**. Scale bar = 50μM

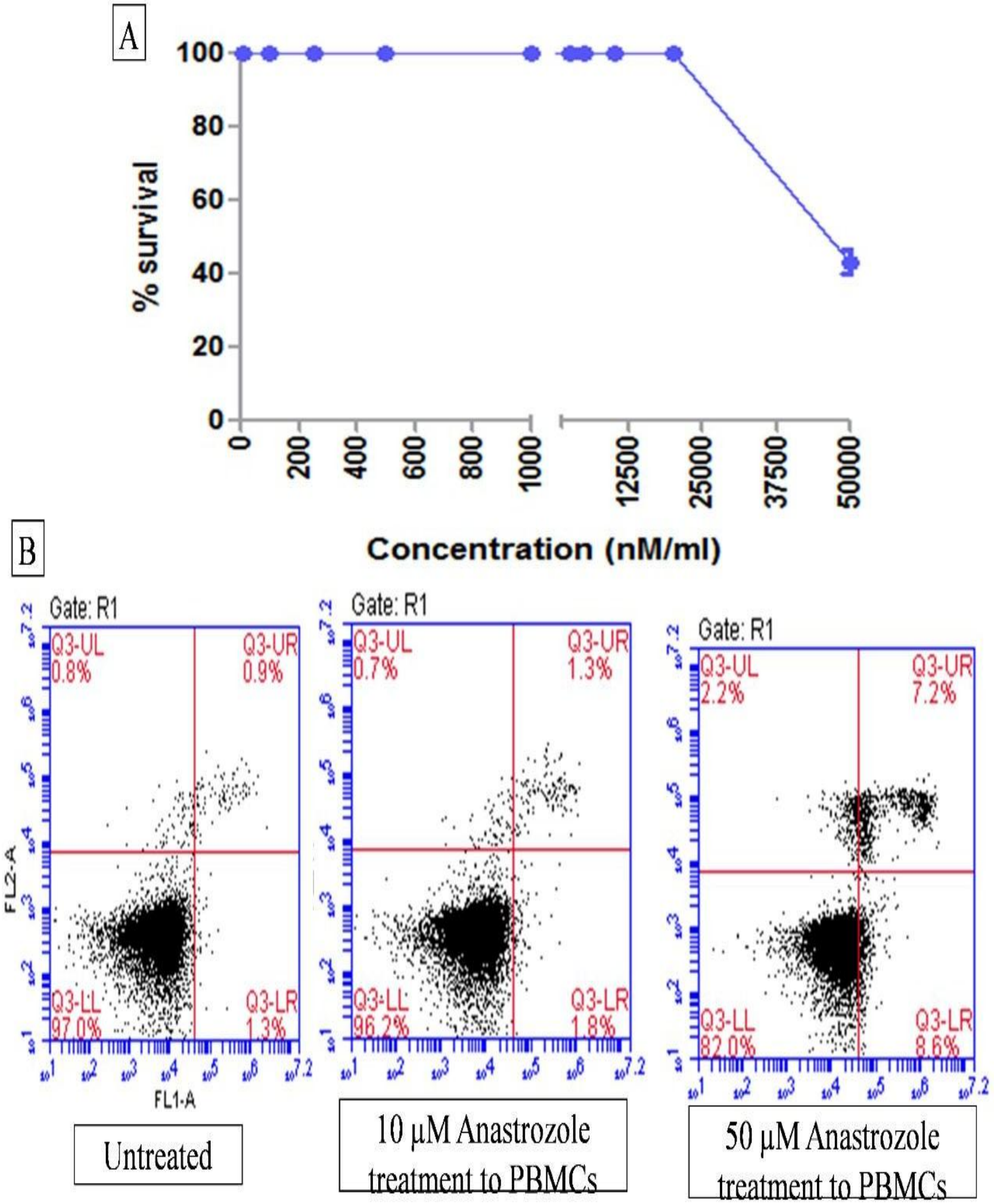


Supplementary Figure-S2 : **Drug toxicity assay**: **(A)** Dosage selection of anastrozole was identified by using the methylthiazol tetrazolium (MTT) assay on human embryonic kidney cells (HEKs). **(B)** Screening of different dosage of anastrozole were treated on HEKs.
